## Supplementary Information for "Emergence of cellular nematic order is a conserved feature of gastrulation in animal embryos"

In the Supplementary Information (SI), we provide additional figures and explanations for the Videos that are pertinent to the results present in the main text.

**SI Figures**

Figure S1. **Snapshots of the mesoderm layer of zebrafish tissues and spatial-temporal evolution of the cell shape index $SI$ during $CE$.**

Figure S2. **Evolution of the cell area distribution in zebrafish.**

Figure S3. **Shape index along the anteroposterior axis at different times.**

Figure S4. **Evolution of the cell shape index outside the notochord.**

Figure S5. **Nematic order during zebrafish development.**

Figure S6. **Spatial-temporal evolution of nematic order during zebrafish development.**

Figure S7. **Evolution of cell orientation during zebrafish $CE$.**

Figure S8. **T1 transitions in zebrafish $CE$.**

Figure S9. **Nucleation and expansion of the nematic phase during zebrafish $CE$.**

Figure S10. **Cell morphology changes during *Xenopus* $CE$.**

Figure S11. **Calculated spatial correlation function $C_S(r)$ (see Eq. 2 in the main text) using Model III**.

Figure S12. **Comparison of cell shape index for wild-type *Xenopus* versus *C-cadherin* knockdown tissues.**

Figure S13. **Calculated nematic order formation using simulations with lattice size 30×30.**

Figure S14. **Computational results using lattice size 50×50.**

**SI Table**

Table I. **The parameters used in the simulation.**

**SI Videos**

Video I. **Movie for zebrafish $CE$.**

Video II. **The evolution of the cell shape index ($SI$) as a function of the cell position ($x$) along the mediolateral axis of zebrafish in Video 1.**

Video III. **The evolution of the cell orientation in zebrafish** *CE***.**

Video IV. **Movie for** *Xenopus laevis CE***.**

Video V. **Movie for** *Drosophila CE***.**

Video VI. **Movie for** *Xenopus laevis CE* **after PCP-protein knockout.**

Video VII. **Movie for** *Xenopus laevis CE* **after Cdh3 knockout.**

Video VIII. **Movie for zebrafish** *CE* **with spt/Tbx16 mutant.**

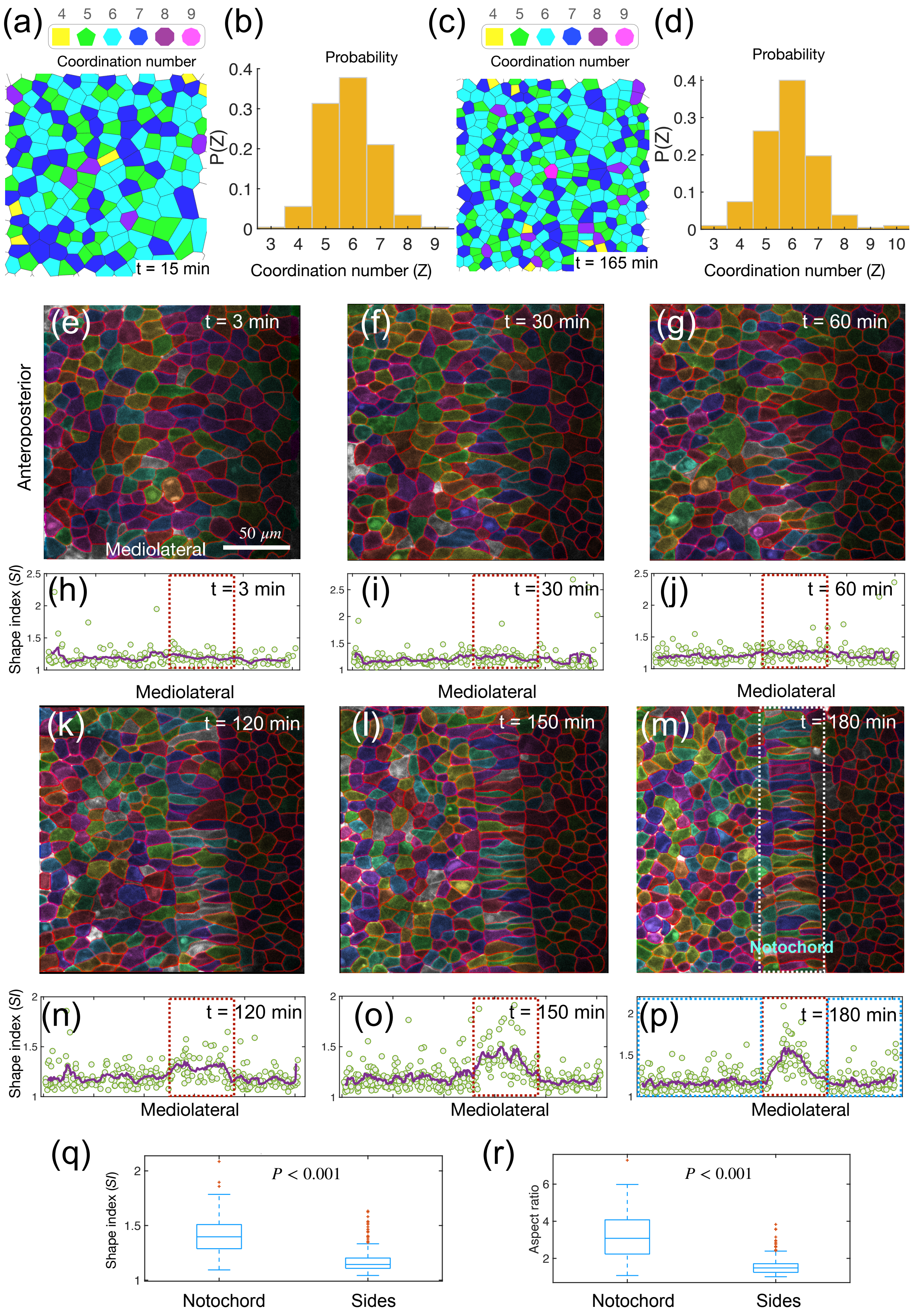
(a)
Coordination number
t = 15 min
(b)
Probability
P(Z)
Coordination number (Z)
(c)
Coordination number
t = 165 min
(d)
Probability
P(Z)
Coordination number (Z)
(e) t = 3 min
Anteroposterior
50 μm
Mediolateral
(f) t = 30 min
(g) t = 60 min
(h) t = 3 min
Shape index (SI)
Mediolateral
(i) t = 30 min
Mediolateral
(j) t = 60 min
Mediolateral
(k) t = 120 min
(l) t = 150 min
(m) t = 180 min
Notochord
(n) t = 120 min
Shape index (SI)
Mediolateral
(o) t = 150 min
Mediolateral
(p) t = 180 min
Mediolateral
(q)
P < 0.001
Shape index (SI)
Notochord
Sides
(r)
P < 0.001
Aspect ratio
Notochord
Sides

FIG. S1: **Snapshots of the mesoderm layer of zebrafish tissues and spatiotemporal evolution of the cell shape index $SI$ during $CE$.** (a), (c) Snapshots of the mesoderm layer of zebrafish at different timepoints, with cells segmented, and cell coordination number Z (the number of neighbors), derived from Voronoi tessellation of the cell centers. Each Voronoi cell is displayed in a different color based on the value of Z, as listed by the polygon symbols at the top of (a), (c). (b), (d) The distribution of Z for cells in (a), and (c). (e)-(g) and (k)-(m) Images of the zebrafish mesoderm layer at different times. Cell colors are for illustration purposes. (h)-(j) and (n)-(p) Shape index, $SI \equiv \frac{P}{\sqrt{4\pi A}}$, as a function of the cell position along the mediolateral axis corresponding to (e)-(g) and (k)-(m). The green dots represent different cells along the anteroposterior axis with a certain x-value (mediolateral axis). The purple line is the average of these points for smoothing the data. (q) Box-plot of the $SI$ of cells located in the notochord region (the red dashed box) versus cells at the two sides (the blue dashed box) in (p). (r) Same as (q), except showing the cell aspect ratio. The two-sided Mann-Whitney U test is used for the statistical analysis. The p-value is listed in (q) and (r).

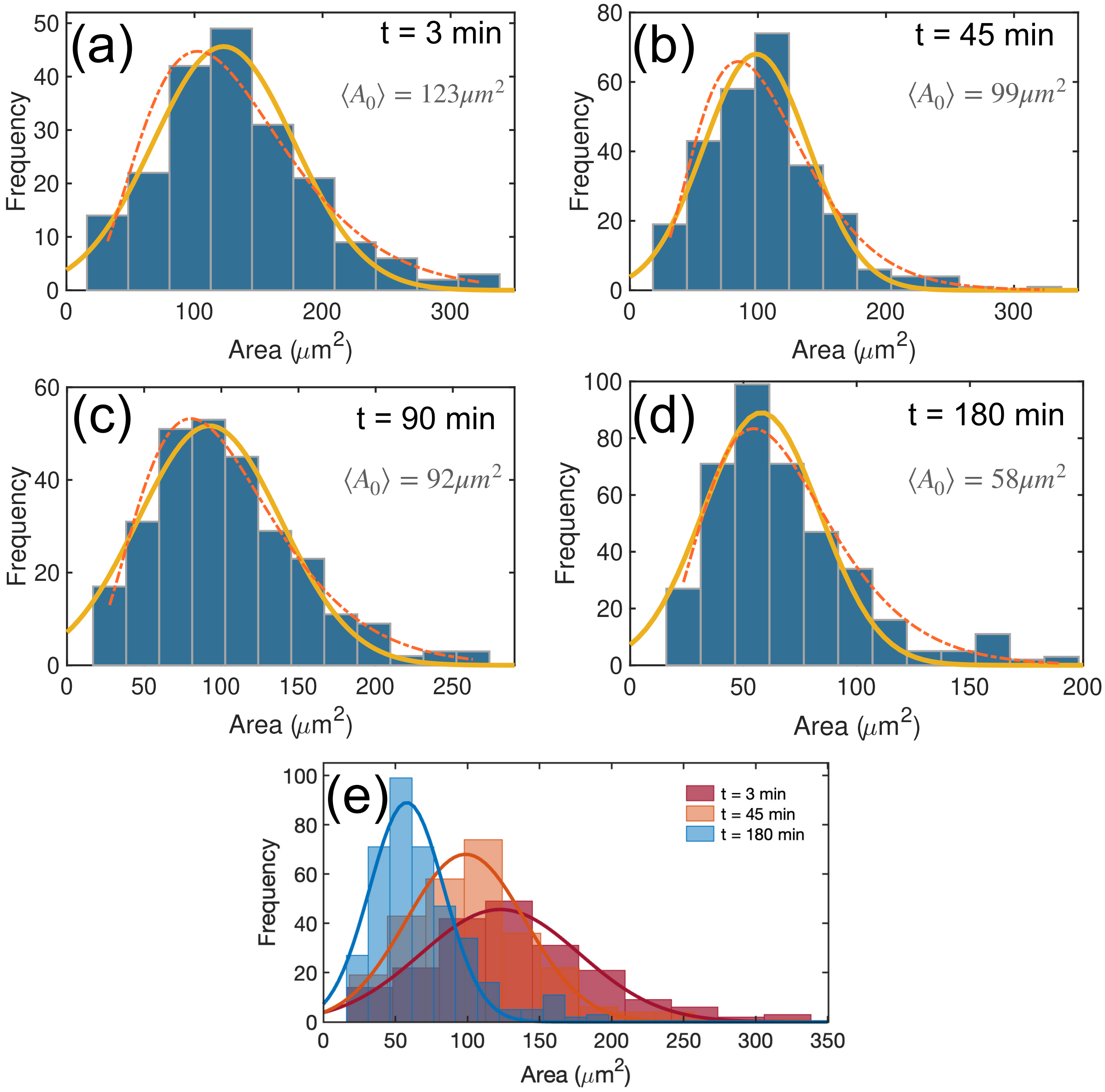


FIG. S2: **Evolution of the cell area distribution in zebrafish.** (a)-(d) Time dependent changes in the cell area distribution. (a)-(d) Histogram of the apical cell area at different times during zebrafish CE. The solid (dash-dotted) line is the Gaussian (Gamma) distribution fit to the data. The mean area, $\langle A_0 \rangle$, of cells from the Gaussian fit is listed in each figure. (e) Distribution of cell area at three time points. There is a shift towards the small values of the mean area as time increases.

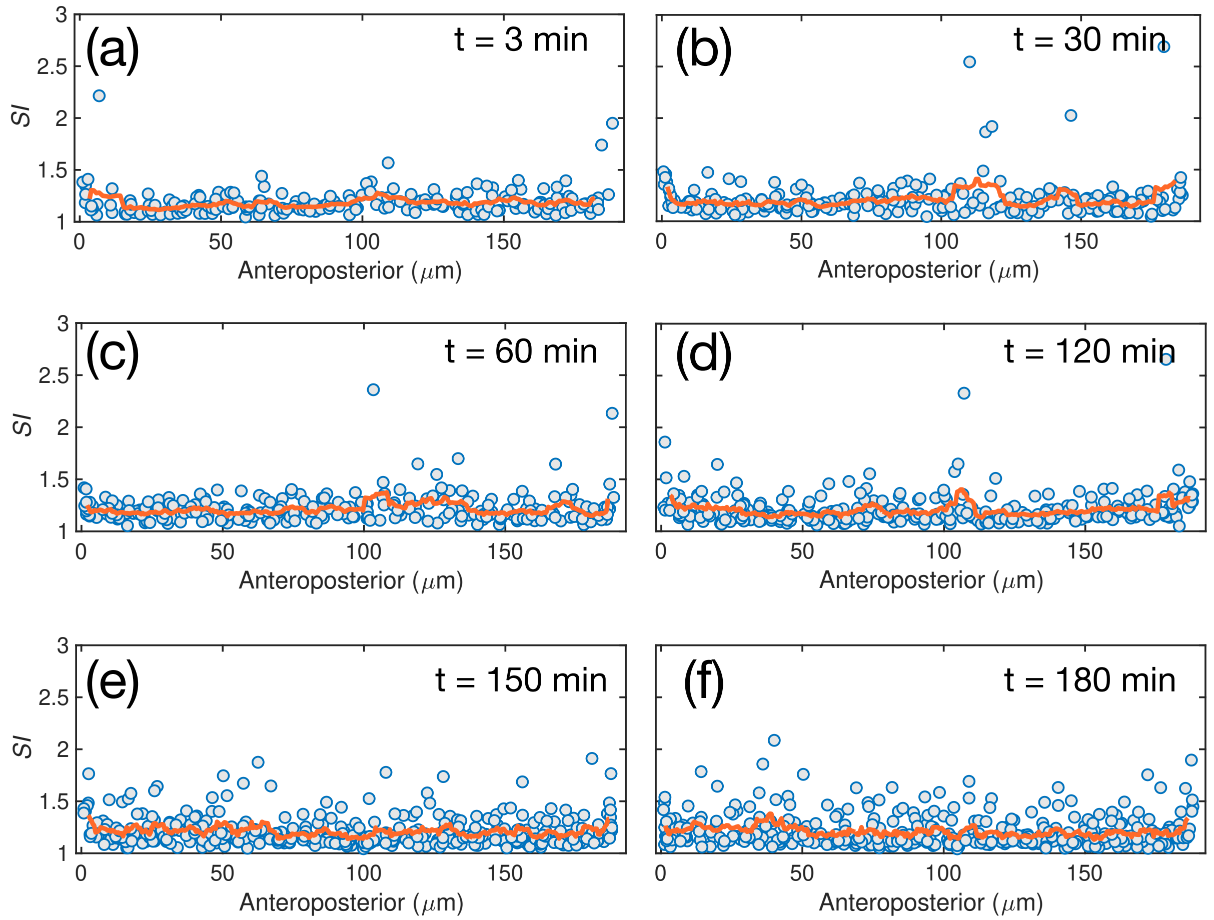


FIG. S3: **Shape index along the anteroposterior axis at different times.** (a)-(f) Same as Figs. S1(h)-(j) and (n)-(p), except for the value of $SI$ as a function of the cell position along the anteroposterior axis corresponding to Figs. S1(e)-(g) and (k)-(m).

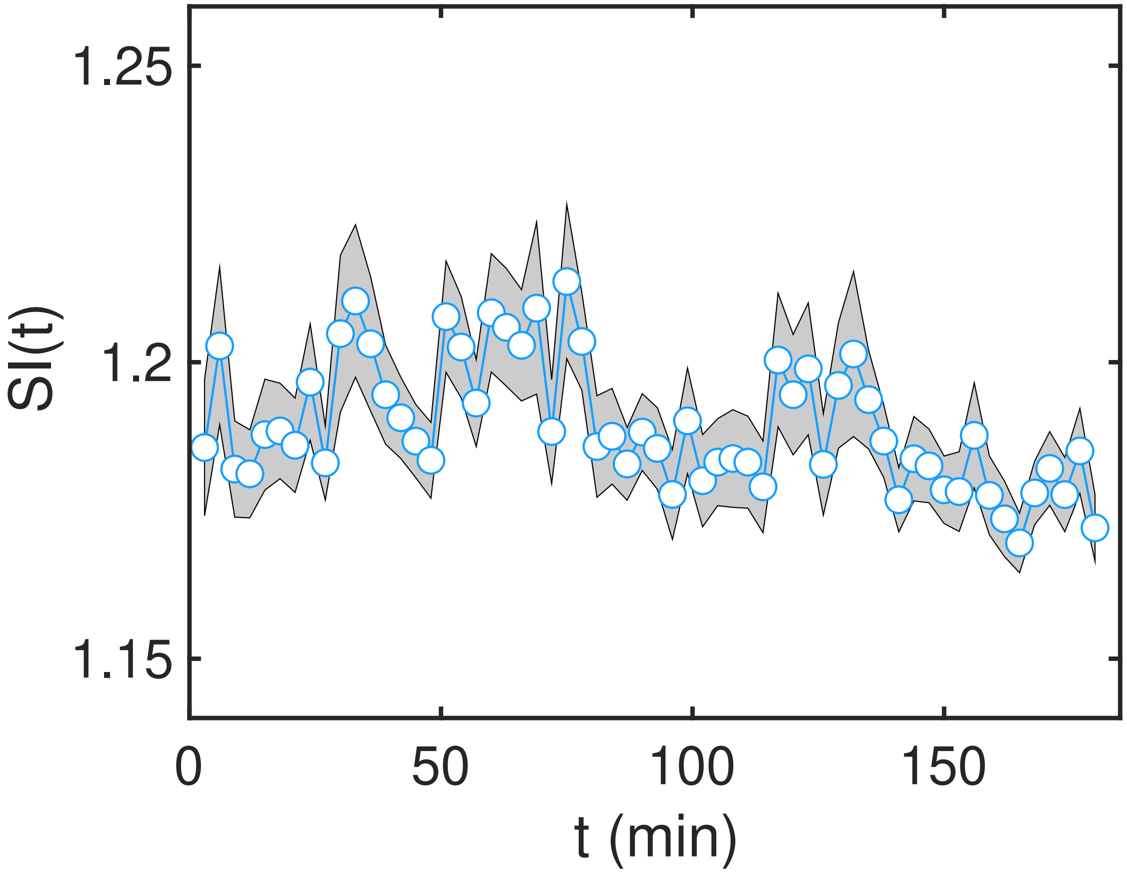


FIG. S4: **Evolution of the cell shape index outside the notochord.** Temporal evolution of the shape index, SI(t), of cells outside the notochord region. No significant change is observed in the time dependence of SI in this region.

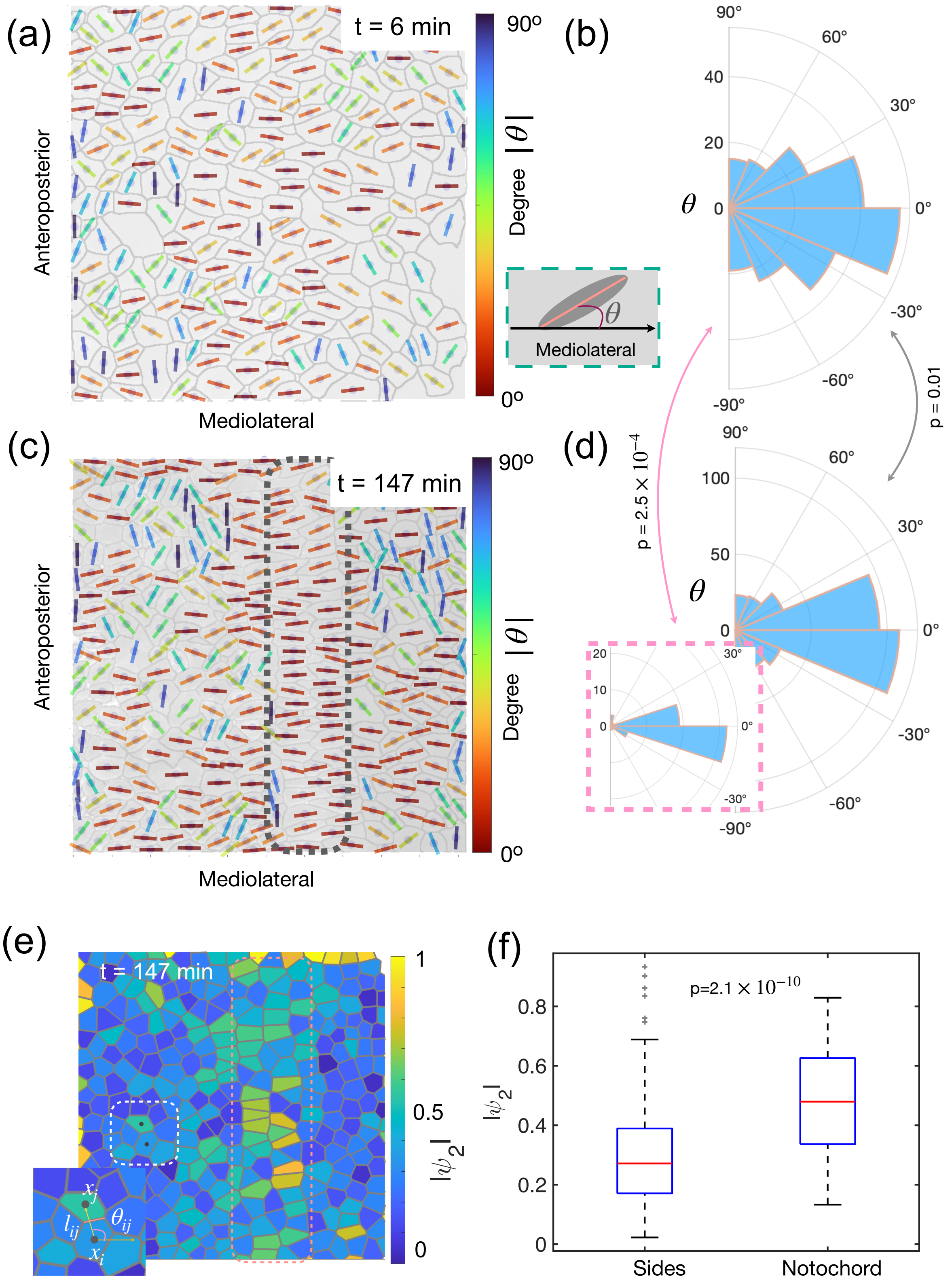


FIG. S5: **Nematic order during zebrafish development.**

FIG. S5: **Nematic order during zebrafish development.** (a) and (c) Same as Figs. 1(d) and (g) in the main text, showing the cell orientation in zebrafish tissues at different times. The cell orientation is defined by the angle, $\theta$, between the long axis of cells (see the short lines) and the horizontal (mediolateral) axis, as shown in the inset at the bottom right of (a). The short lines are color coded by the value of $|\theta|$. (b) Histogram of the polar angle $\theta$ for all cells in (a). (c)-(d) Same as (a)-(b), except they are at a later timepoint. The inset in (d) gives the histogram of $\theta$ for cells located in the notochord region shown by the pink dotted rectangle in (c). The p-values between the patterns in (b) and (d), (b) and the inset in (d) are calculated with two-sample Kolmogorov–Smirnov test. (e) The 2-fold orientation order, $|\psi_2|$, of cells (see the definition in Eq.(3) in the Materials and Methods section). Each Voronoi cell is color coded by the value of $|\psi_2|$ (see the scale on the right). The inset on the left bottom is a zoom in of the white dashed rectangle region. The variables $\theta_{ij}$ (the angle between the vector pointing from cell i to j and the horizontal axis) and $l_{ij}$ (the length of the edge shared between the Voronoi cells i and j) are used in the definition of $|\psi_2|$. (f) Boxplot of $|\psi_2|$ for cells located at two sides or the middle region (see the dotted rectangle in (e)). The p-value is calculated using the two-sided Mann-Whitney U test.

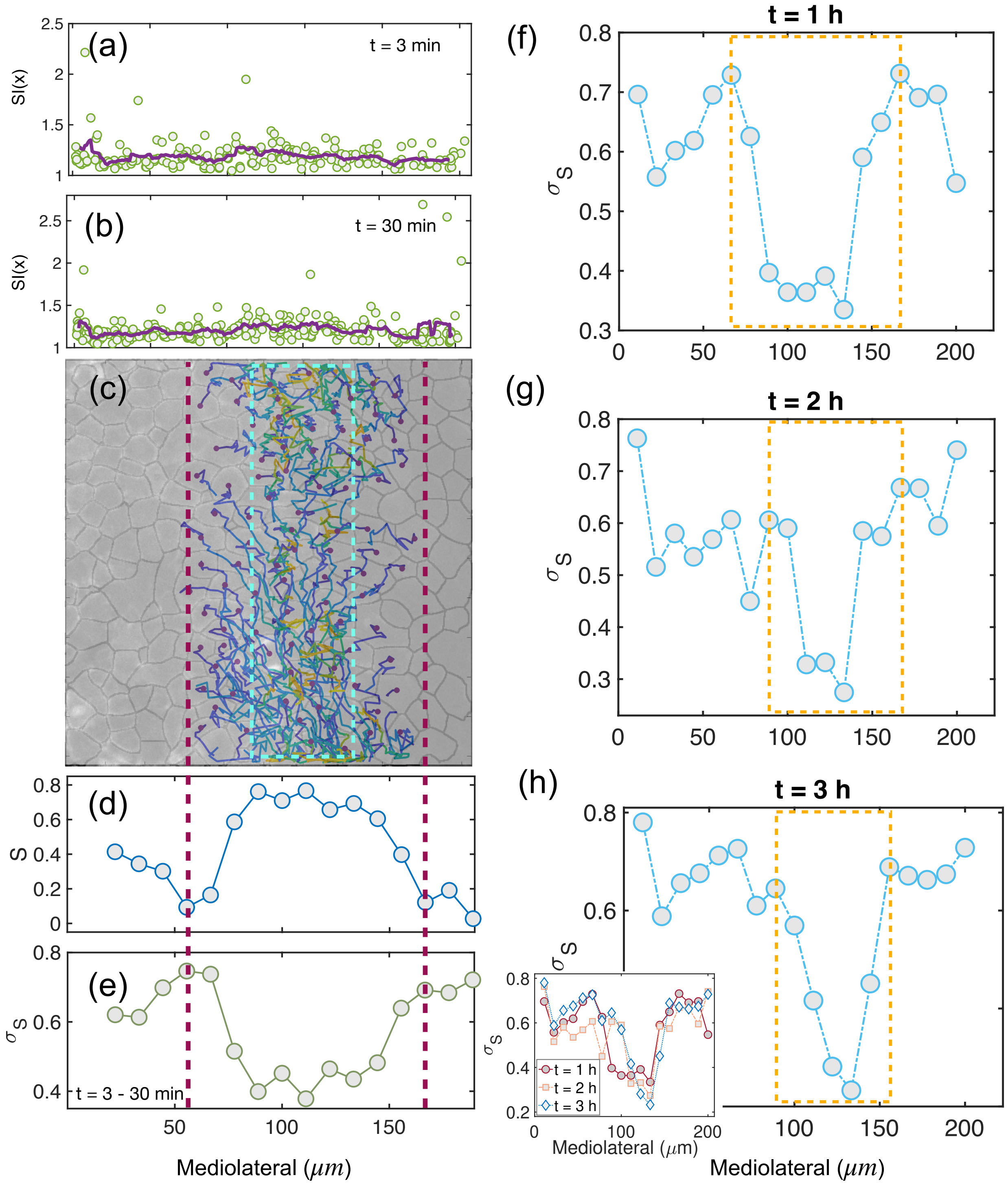


FIG. S6: **Spatiotemporal evolution of nematic order during zebrafish development.**

FIG. S6: **Spatio-temporal evolution of nematic order during zebrafish development.** (a)-(b) The cell shape index, $SI(x)$, along the mediolateral axis at early times. (c) The tissue snapshot at an early time point is overlaid with the trajectories of cells located inside of the red dashed lines over 3 hours, with blue color for early time points and yellow for late timepoints. The dashed rectangle in cyan gives the final notochord region. (d)-(e) The nematic order parameter, S, and the standard deviation, $\sigma_S$, along the mediolateral axis. The value of S and $\sigma_S$ in (d)-(e) are averaged over 10 successive time frames (t=30 minutes). Note the fluctuations in the notochord is significantly less than for cells that are outside it. The boundaries between the nematic-order and disordered regions are indicated by the red dashed lines. (f)-(h) The spatial-temporal evolution of $\sigma_S$. A sharp transition of $\sigma_S$ along the mediolateral axis is observed as indicated by the dashed rectangles at different times, and this region narrows over time. The inset in (h) shows the value of $\sigma_S$ at different times.

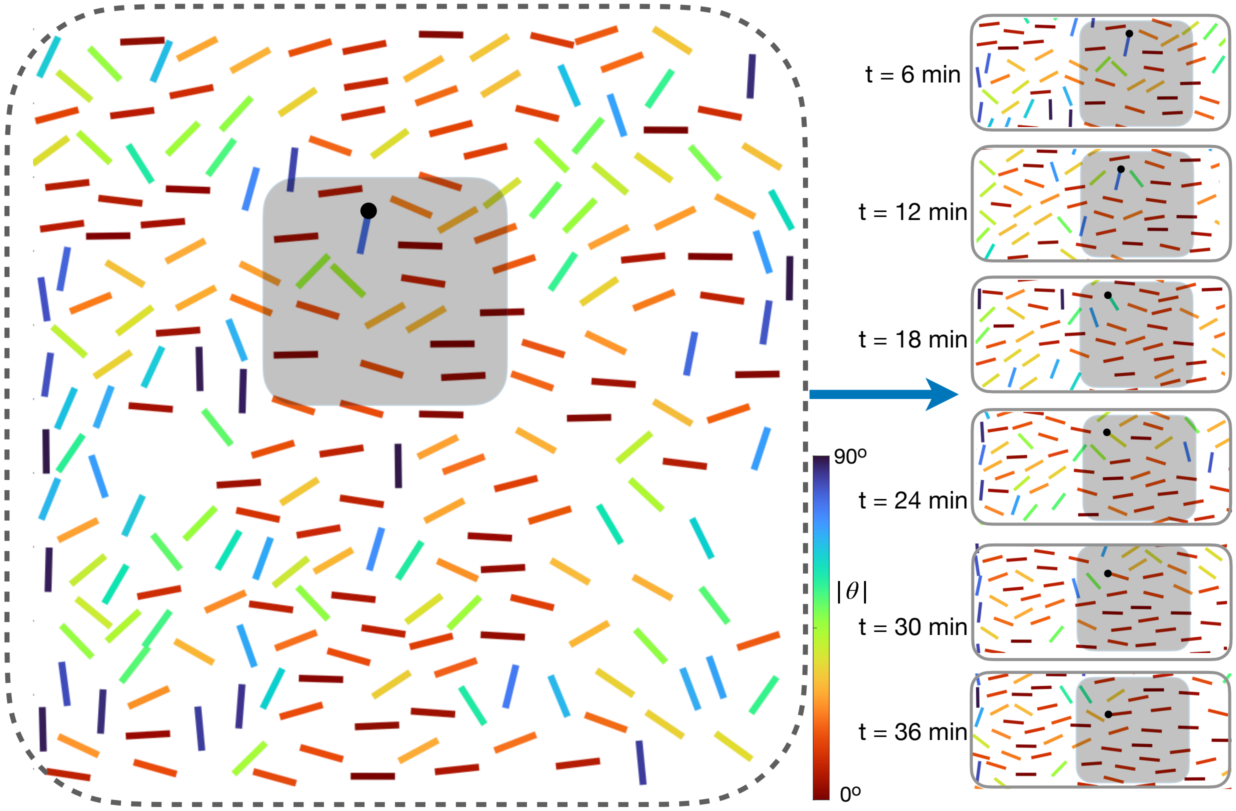


FIG. S7: **Evolution of cell orientation during zebrafish** ***CE*****.** Similar to Fig. 1(d) in the main text. The cell orientation is defined by the angle, $\theta$, between the long axis of cells (see the short lines) and the mediolateral (horizontal) axis as shown in the inset in Fig. S5(a). The lines are color coded by the value of $|\theta|$. Figures on the right show the evolution of cell orientation located in the shaded area of the left figure. There are dramatic changes in the orientation of individual (see the line with a dot on one side as an example) in the process of forming a nematic phase.

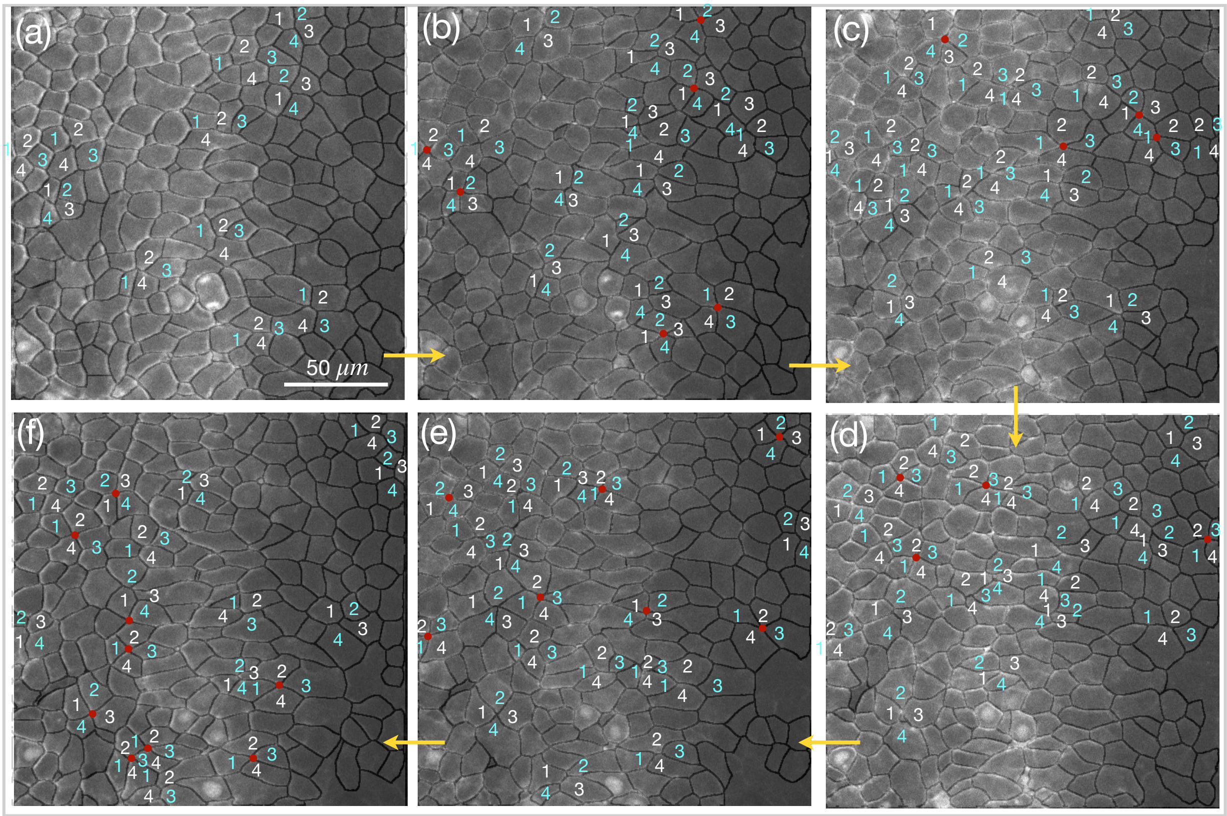


FIG. S8: **T1 transitions in zebrafish** ***CE*****.** Several examples of T1 transitions in zebrafish *CE* in six consecutive images. Four cells are labeled 1, 2, 3 and 4 with white color indicating two cells that are in contact with each other while cyan color indicates they are separated. The red dot shows where two vertices meet.

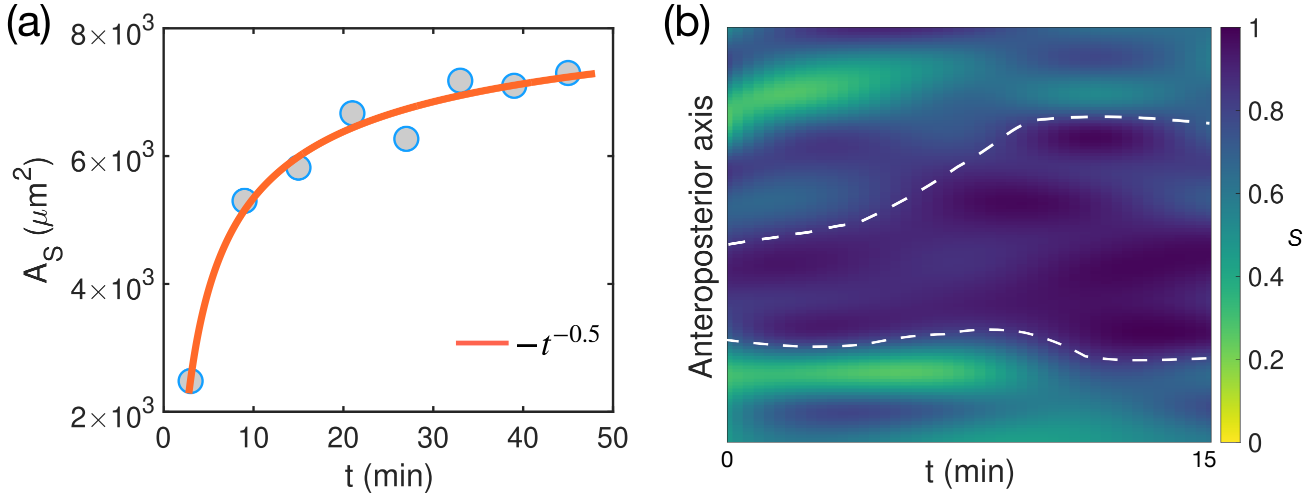


FIG. S9: **Nucleation and expansion of the nematic phase during zebrafish** ***CE*****.** (a) The area of the nematic phase, $A_S$ (with $S > 0.8$), as a function of time in the notochord region corresponding the Fig. 1l in the main text. (b) A kymograph showing the mean values of S (averaged over the mediolateral axis on the notochord region) along the anteroposterior axis at different times. The dashed lines (indicating the boundaries of the nematic order region) are a guide to the eye, showing the nucleation and growth of the nematic order.

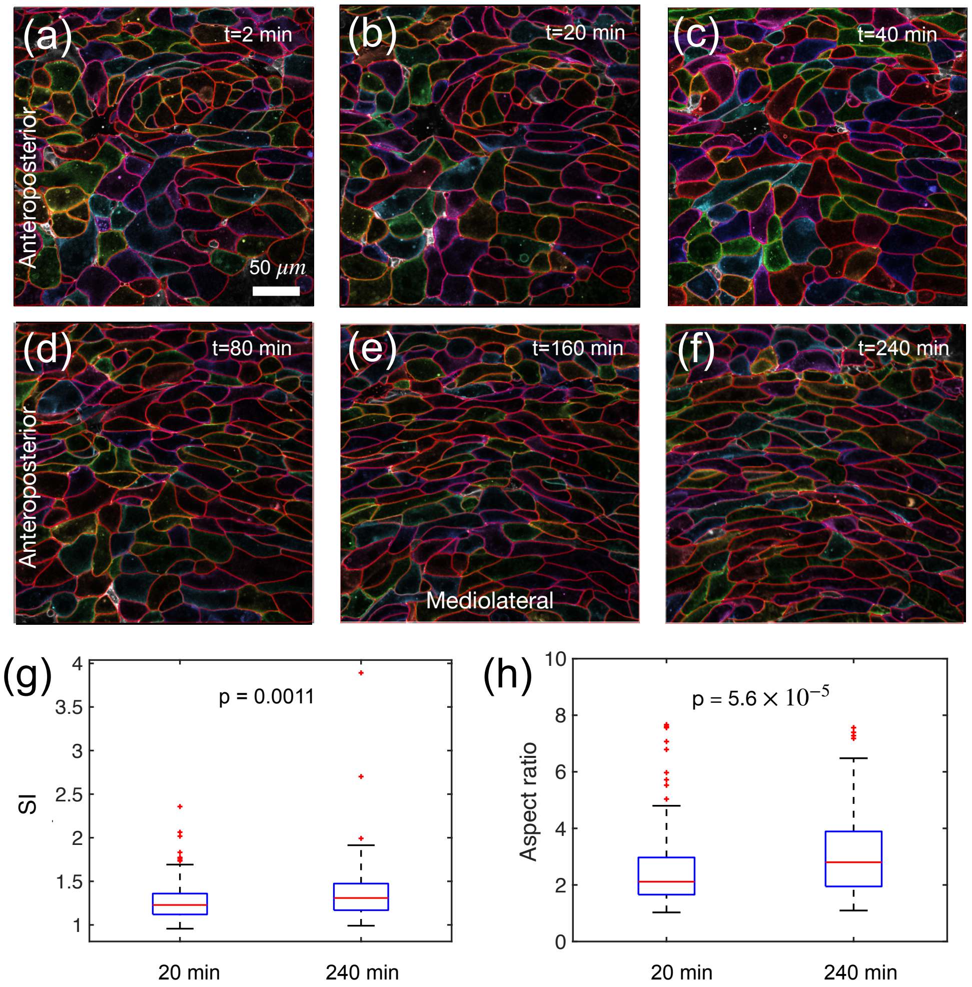


FIG. S10: **Cell morphology changes during** ***Xenopus CE*****.** (a)-(f) Snapshots of *Xenopus laevis* tissues at different times during *CE*. (g) Box-plot of the *SI* value calculated for cells from (b) and (f). (h) Aspect ratio of cells from (b) and (f). The two-sided Mann-Whitney U test is used for the statistical analysis in (g)-(h). The p-value is listed in each figure.

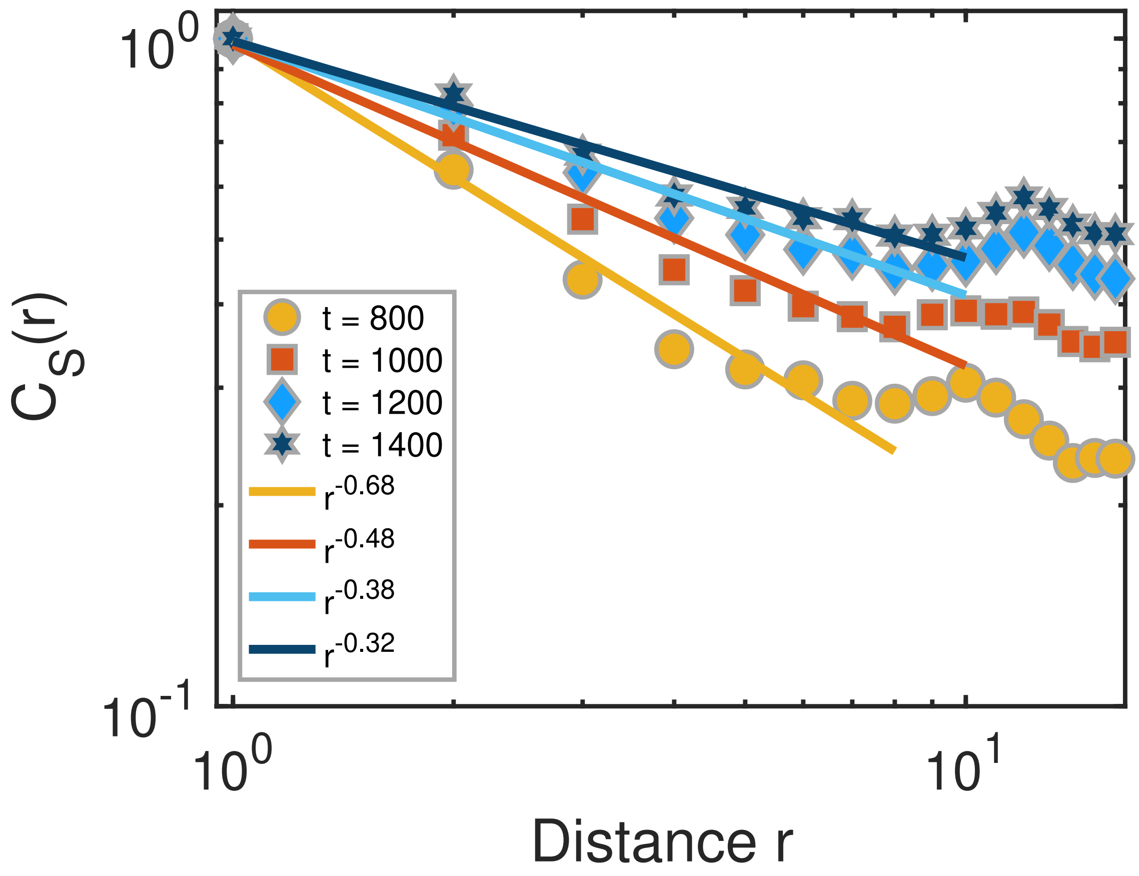


FIG. S11: **Calculated spatial correlation function** $C_S(r)$ **(see Eq. 2 in the main text) using Model III**. The spatial correlation, $C_S(r)$, of cells at different times from simulations in Model III with the same parameters used in Fig. 4i-l in the main text. The solid lines are the power-law fits to the data.

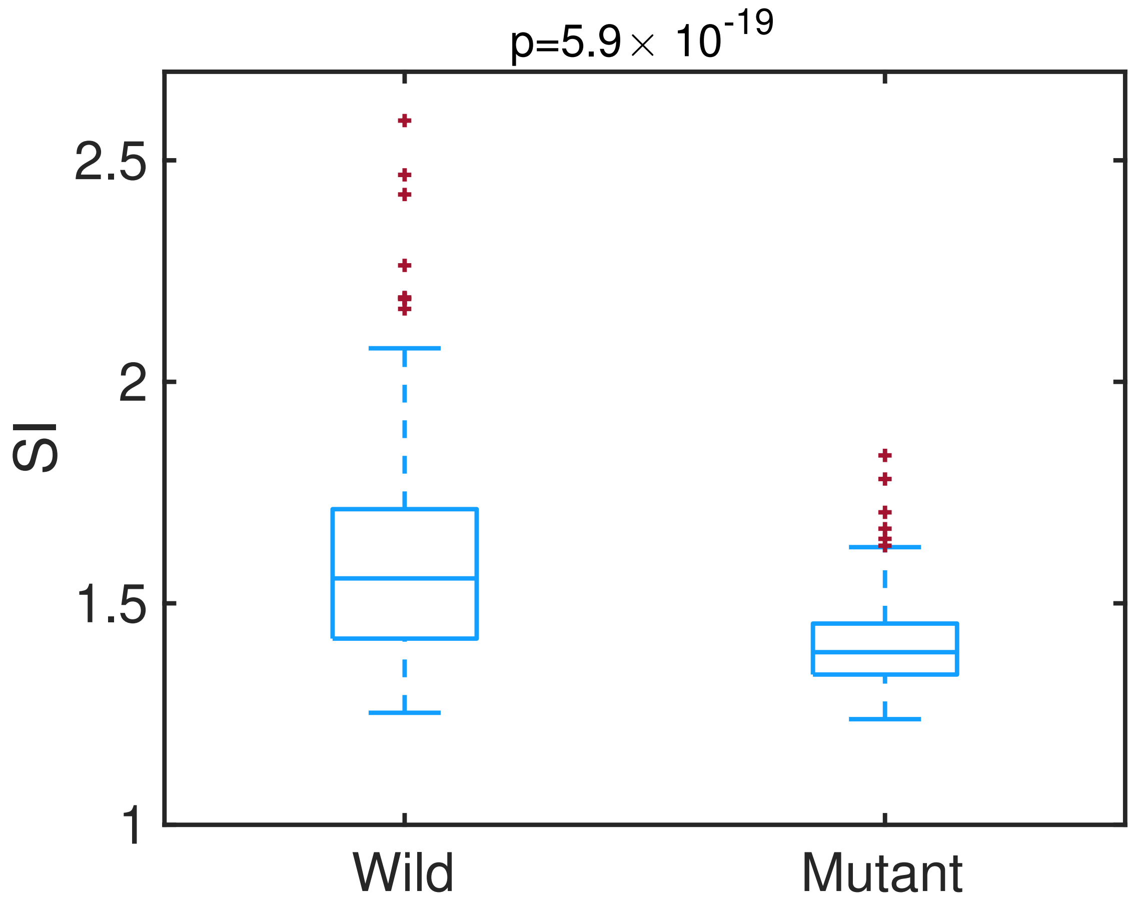


FIG. S12: **Comparison of cell shape index for wild-type *Xenopus* versus *C-cadherin* knockdown tissues.** Box-plot of the $SI$ for cells from Figs. 6(a) and (b) in the main text for *C-cadherin* knockdown and wild-type *Xenopus* tissues. Mann-Whitney U test is used for the statistical analysis. The p-value is listed in the figure.

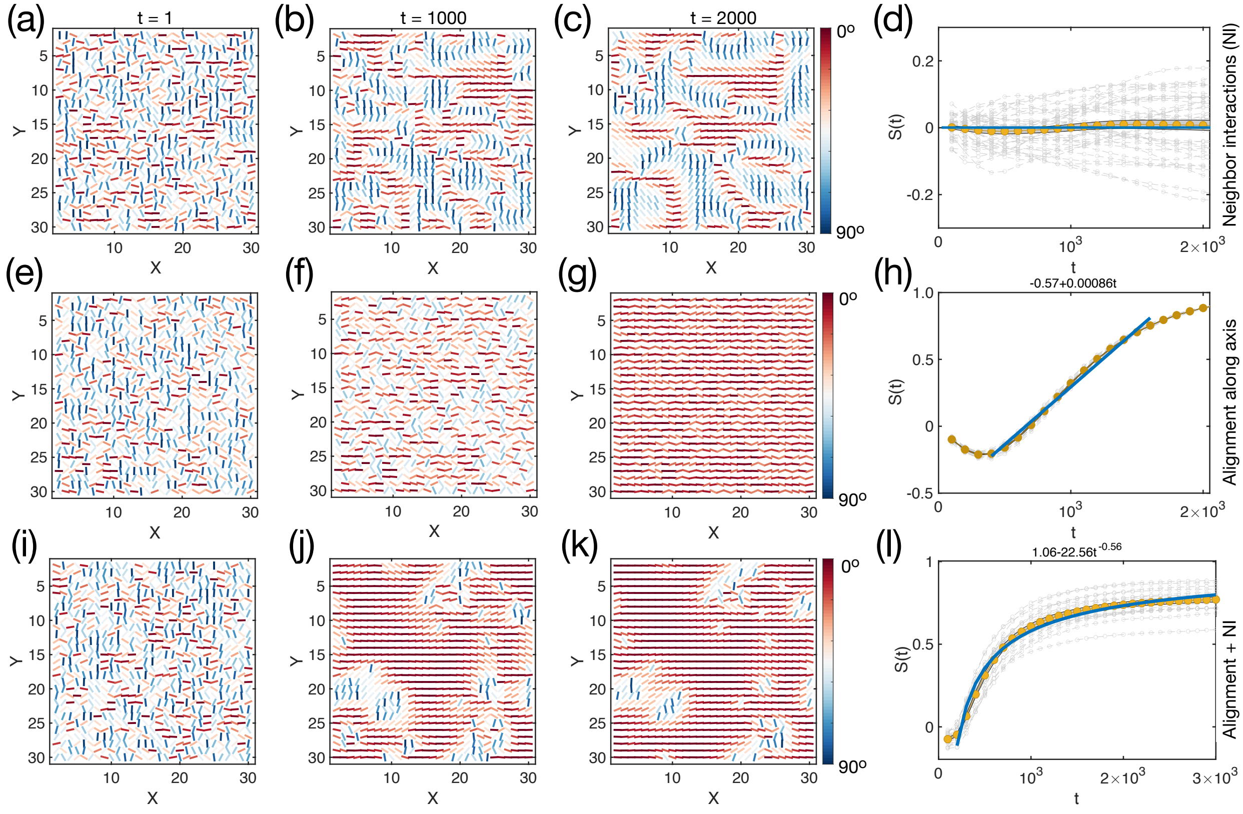


FIG. S13: **Calculated nematic order formation using simulations with lattice size 30×30.** Same as Figure 4 in the main text, except the lattice size is 30×30. (a)-(d) Results from model (i). (e)-(h) Results from model (ii). (i)-(l) Results from model (iii). The parameter $\mathcal{A}$=$2.5\times10^{-3}$, $\mathcal{B}$=$1\times10^{-3}$. The models are described in the main text.

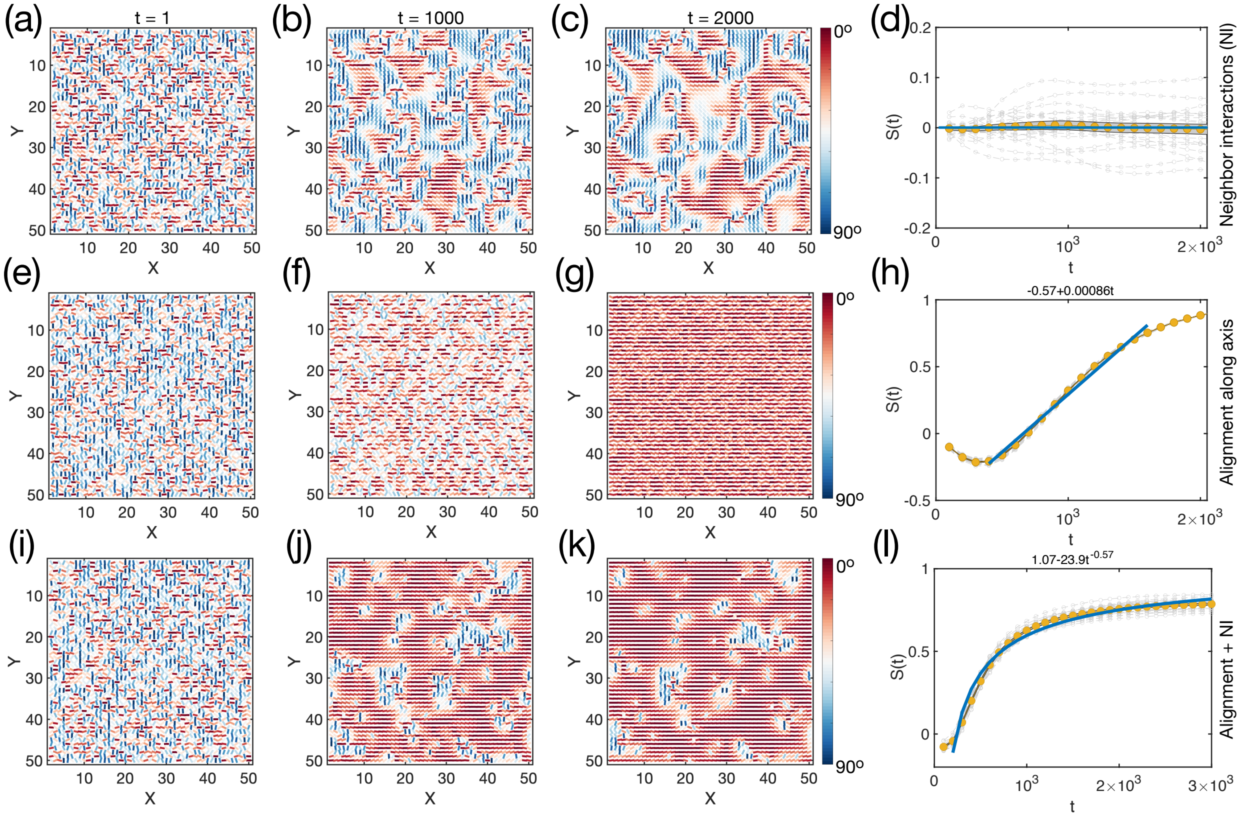


FIG. S14: **Computational results using lattice size 50×50.** Same as Figure S13, except the lattice size is 50×50.

TABLE I: The parameters used in the simulation.

| **Models** | $\mathcal{A}(local)$ | $\mathcal{B}(global)$ |
|---|---|---|
| Model i | $2.5\times10^{-3}$ | 0 |
| Model ii | 0 | $1\times10^{-3}$ |
| Model iii | $2.5\times10^{-3}$ | $1\times10^{-3}$ |
| Model iii (*Xenopus* mutant) | $10^{-6}$ | $3\times10^{-4}$ |
| Model iii (zebrafish mutant) | $2.5\times10^{-3}$ | $-1\times10^{-3}$ |
